# Commensal *Escherichia coli* colonization triggers Peyer’s patch development

**DOI:** 10.1101/2025.10.06.680803

**Authors:** Romana R. Gerner, Gregory T. Walker, Suzi M. Klaus, Karine Melchior, Suzana Hossain, Kareem Siada, Chia-Yun Hsu, Francisco J. Albicoro, William Santus, Rasika Patkar, Marvic Carrillo-Terrazas, Grant J. Norton, Flavian Thelen, Araceli Perez-Lopez, Purnima Sharma, Marcus P. Wong, Victor Lei, Richard M. Ransohoff, David D. Lo, Thomas E. Lane, Andrea Reboldi, Sean-Paul Nuccio, Judith Behnsen, Elina I. Zuniga, Li-Fan Lu, Çagla Tükel, Hiutung Chu, Manuela Raffatellu

**Affiliations:** Division of Host-Microbe Systems and Therapeutics, Department of Pediatrics, University of California San Diego, La Jolla, CA 92093, USA; School of Life Sciences, Freising-Weihenstephan, ZIEL – Institute for Food & Health, Freising-Weihenstephan, Technical University of Munich, Germany; Department of Internal Medicine III, School of Medicine, University Hospital rechts der Isar, Technical University of Munich, Munich, Germany; Department of Microbiology and Molecular Genetics, University of California, Irvine School of Medicine, Irvine, CA 92697, USA; Department of Pathology, University of California, San Diego, La Jolla, CA, USA; Department of Microbiology and Immunology, Lewis Katz School of Medicine, Temple University, Philadelphia, PA, USA; Department of Microbiology and Immunology, University of Illinois Chicago, Chicago, IL 60612, USA; Department of Biological Sciences, University of Illinois Chicago, Chicago, IL, 60612, USA; School of Biological Sciences, University of California, San Diego, La Jolla, California, USA; Third Rock Ventures, Boston, MA 02215; Division of Biomedical Sciences, University of California, Riverside, School of Medicine 900 University Ave, Riverside, CA 92521; Department of Neurobiology and Behavior, University of California, Irvine 92697-3900, USA; Department of Pathology, UMass Chan Medical School, Worcester, MA, USA; Chiba University-UC San Diego Center for Mucosal Immunology, Allergy, and Vaccines (CU-UCSD cMAV), La Jolla, CA, 92093, USA; Center for Microbiome Innovation, University of California, San Diego, La Jolla, CA 92093, USA

## Abstract

The gut microbiota plays a pivotal role in shaping mucosal immunity, yet the specific microbes contributing to lymphoid tissue development remain poorly defined. Here, we identify *Escherichia coli*, a pioneer commensal bacterium, as a key driver of naïve B cell accumulation in gut Peyer’s patches and lamina propria via a CXCR2-dependent mechanism. We show that *E. coli* promotes B cell recruitment through the production of curli amyloid fibers, which signal via Toll-like receptor 2 (TLR2). Notably, this effect extends beyond the neonatal period, revealing a broader temporal window for microbial modulation of mucosal immune development. These findings reveal a previously unrecognized role for a defined gut commensal bacterium and its molecular products in orchestrating the formation of gut-associated lymphoid tissue and B cell recruitment.

## Main Text

The gastrointestinal tract harbors a highly complex and diverse microbial community, known as the gut microbiota, which plays fundamental roles in the maturation of gut-associated lymphoid tissue (GALT) (*1*, *2*). The intestine contains the largest population of immune cells in the human body, fostering continuous and dynamic interactions between the immune system and the microbiota. Over the past 15 years, seminal studies have significantly advanced our understanding of how specific gut microbes shape immune development (*3*). For example, segmented filamentous bacteria (SFB) promote the differentiation of T helper 17 cells via epithelial cell-mediated endocytosis of microbial antigens (*4*, *5*). Similarly, *Bacteroides fragilis* modulates the development of regulatory T cells via the secretion of a capsular polysaccharide (*6*). Still, the influence of specific microbiota members on immune cell trafficking and the underlying mechanisms for the development of gut lymphoid tissues remains largely elusive, highlighting a significant gap in our understanding of mucosal immunology.

Exposure to the gut microbiota influences the B cell repertoire and the production of antibodies, particularly immunoglobulin A (IgA) (*7–9*). B cells are distributed throughout the gastrointestinal tract and are particularly enriched within organized lymphoid structures. The most prominent of these are Peyer’s patches: dome-shaped lymphoid follicles located along the small intestine. Peyer’s patches are lined by specialized epithelial cells known as microfold cells (M cells), which transport bacteria and luminal antigens toward the underlying immune cell aggregates. These processes trigger protective immunity or induce tolerance toward commensal organisms, with Toll-like receptor (TLR) signaling playing a central role in many of these pathways (*10*, *11*). Although Peyer’s patch development is initiated prenatally during gestation, the postnatal recruitment and maturation of immune cells within these structures are primarily driven by microbial exposure (*12–14*). Indeed, early-life antibiotic exposure disrupts Peyer’s patch development (*15*). Additionally, germ-free mice have underdeveloped and smaller GALT structures, including Peyer’s patches, compared to conventional mice (*16*, *17*), and Peyer’s patch development occurs after colonization with a complex microbiota (*18*). Although microbial exposure contributes to postnatal Peyer’s patch development, the roles of specific microbes and the underlying mechanisms remain to be identified.

By comparing mice from different colonies, germ-free mice, and monocolonized mice, we discovered that commensal *Escherichia coli* is a potent inducer of Peyer’s patch development and B cell accumulation in the GALT in both adult and neonate mice. Mechanistically, we found that the production of the *E. coli* amyloid curli and signaling via TLR2 are essential for this process. Colonization with *E. coli* induces the expression of CXC chemokines in Peyer’s patches, and signaling through the chemokine receptor CXCR2 promotes B cell accumulation in the GALT. Our study thus highlights a key role of curli-producing commensal *E. coli* in the development of mucosal immunity.

## Results

### Peyer’s patch size differs in adult mice from different vendors

Microbial colonization promotes GALT development in germ-free mice, including the accumulation and differentiation of lymphocytes in Peyer’s patches (*16*). When comparing mice from different commercial vendors, we noticed substantial variability in Peyer’s patch size among specific pathogen-free (SPF) mice from different vendors. Adult mice from The Jackson Laboratory (Jackson) have small, underdeveloped Peyer’s patches, comparable in size to those observed in germ-free mice. In contrast, mice from Taconic Biosciences (Taconic) harbor significantly larger Peyer’s patches distributed along the small intestine (**Fig. 1, A and B**). The total number of Peyer’s patch B and T cells was also significantly lower in germ-free and Jackson mice compared to Taconic mice (**Fig. 1C**). These observations prompted us to hypothesize that microbial colonization *per se* is insufficient to promote postnatal Peyer’s patch development and that specific microbes may be involved in this process.

**Fig 1.**
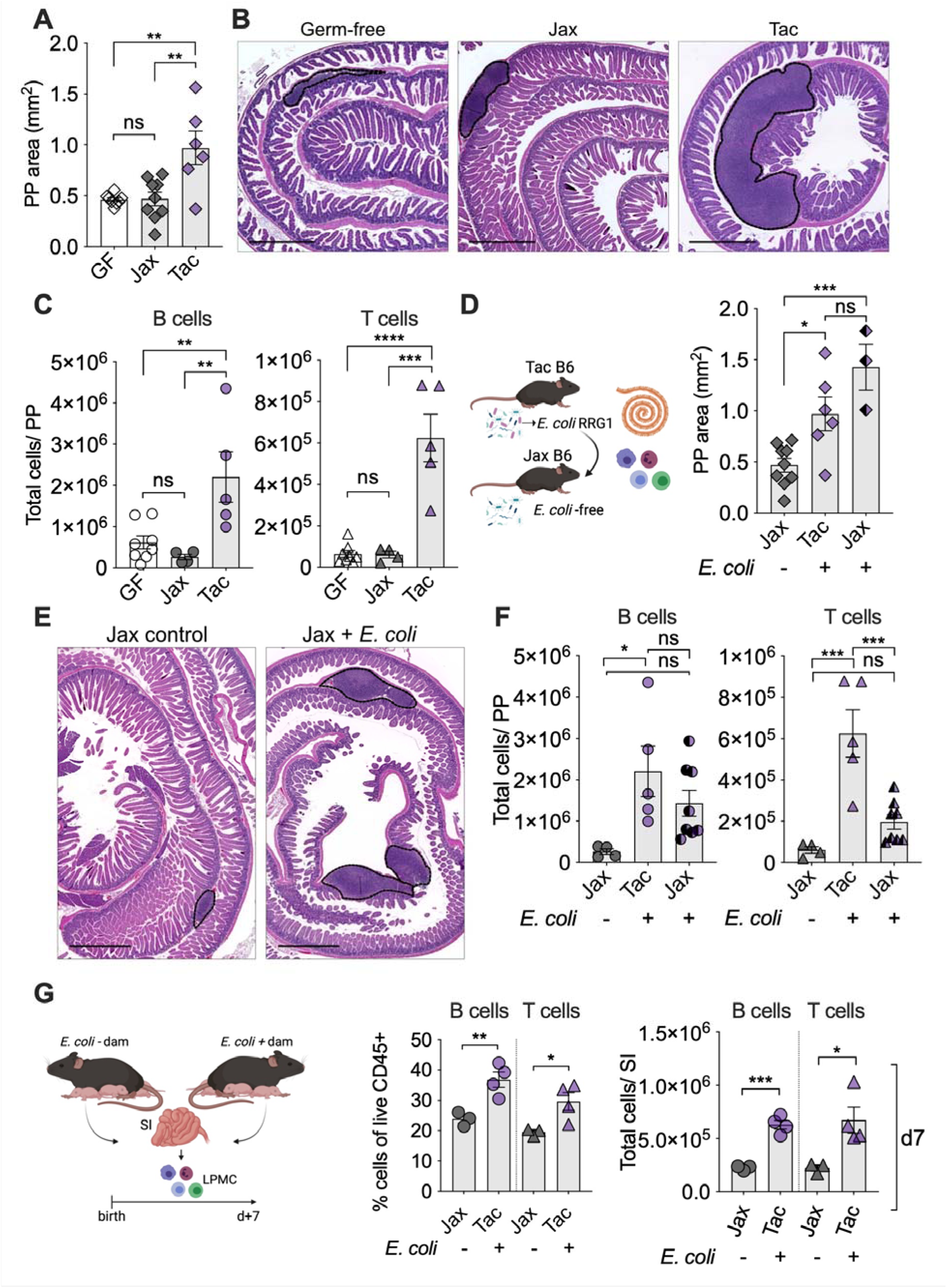
*E. coli* promotes Peyer’s patch development in adult mice. **(A)** Quantitative analysis of total Peyer’s patch (PP) areas per mouse based on H&E-stained Swiss roll sections of the small intestine. Comparisons were made between germ-free (GF) mice and specific pathogen-free (SPF) mice obtained from Jackson Laboratories (Jax) and Taconic Farms (Tac). (**B**) Representative H&E-stained Swiss roll sections of the small intestine from GF, Jax, and Tac mice are shown. Dashed black lines delineate PP areas. 50X magnification, scale bars 1mm. (**C**) Absolute numbers of B (CD45^+^CD19^+^B220^+^) and T (CD45^+^CD3^+^) cells in all PP per small intestine of GF, Jax, and Tac mice. (**D** and **E**) Experiment outline. Jax mice were either colonized with *E. coli* RRG1 for 7 days or uninoculated as a control. Swiss roll sections of the small intestine were then prepared for analysis of PP areas. Dashed black lines delineate PP areas. 50X magnification, scale bars 1 mm. (**F**) In separate experiments, all PP per mouse from the small intestine of Jackson and Taconic mice were excised, and B and T cells were quantified by flow cytometry. (**G**) B and T cells from the small intestines of 7-day-old pups born to Jax dams (*E. coli*-negative) or Tac dams (*E. coli*-positive) were quantified by flow cytometry. Each symbol represents an individual mouse. LPMC, lamina propria mononuclear cells. To minimize the use of experimental animals, data points from panels 1A and 1C (Jax mice without *E. coli* & Tac mice) were reused in panels 1D and 1F, respectively. Bars represent the mean ± SEM. Significant differences are indicated by *p ≤0.05, **p<0.01, ***p<0.001, ****p<0.0001, ns = not significant. One-way ANOVA followed by Tukey’s multiple-comparison test. Illustrations were created using BioRender.

### *E. coli* promotes Peyer’s patch development in adult mice

Significant differences exist in the microbiota composition of Jackson and Taconic mice. Taconic but not Jackson mice are colonized with SFB, which is crucial for the induction of Th17 cells (*4*). Taconic mice are also colonized with commensal *Enterobacteriaceae*, which promote resistance to infection with enteric pathogens (*19*). In contrast, Jackson mice do not harbor *Enterobacteriaceae* in their microbiota (*19*). Consistent with this earlier study, cultured fecal samples from Taconic mice, but not Jackson mice, showed growth of *E. coli*, as evidenced by lactose-fermenting colonies on MacConkey plates and confirmed by whole-genome sequencing (**fig. S1, A and B**). *E. coli* is an important member of the gut commensal microbiota of mammals, yet its relative abundance is low in a healthy gut (*20*). To test whether the difference in *E. coli* colonization could explain the differences in Peyer’s patch size, we administered the commensal *E. coli* strain isolated from Taconic mice (designated RRG1) to Jackson mice by oral gavage one day after streptomycin treatment to ensure maximal colonization. Strikingly, the total Peyer’s patch areas of Jackson mice colonized with *E. coli* increased significantly and were comparable to those of Taconic mice after 7 days. Moreover, Jackson mice colonized with *E. coli* showed a significant increase in B cell numbers in the Peyer’s patches, whereas T cell numbers were only partially restored (**Fig. 1, D to F, and fig. S1C**).

### *E. coli* colonization enhances B cell trafficking to the gut in early life

Microbial colonization in early life is critical for GALT development (*13–15*, *21*). While *Enterobacteriaceae* comprise a small fraction of the gut microbiota in healthy adults, they are predominant in early life, particularly in newborns (*22*, *23*), when oxygen levels are higher (*24*). To investigate whether *E. coli* colonization triggers intestinal B cell accumulation during the early postnatal period in mice, we compared neonates born to *E. coli*+ (Taconic) or *E. coli*- (Jackson) dams. As the Peyer’s patches are not macroscopically visible in the neonatal mouse gut, we analyzed lamina propria B and T cells from the small intestine by flow cytometry on day 7 after birth (**Fig. 1G**). Consistent with our prediction, we found a significantly higher frequency and total number of CD45^+^CD19^+^B220^+^ B cells and CD45^+^CD3^+^ T cells in the lamina propria of pups born from *E. coli*+ dams. Thus, early-life *E. coli* colonization contributes to B and T cell recruitment to the gut.

### *E. coli* colonization triggers Peyer’s patch enlargement in germ-free mice

We next investigated whether monocolonization with representative members of the human gut microbiota would promote Peyer’s patch development. We gavaged germ-free mice with commensal model organisms, followed by histological analysis of Peyer’s patches on Swiss roll sections after seven days. Commensals typically found in the human gut, including *Bacteroides thetaiotaomicron*, *Clostridium* sp. 7_2, *Enterobacter cloacae*, *Proteus mirabilis*, and *Lacticaseibacillus rhamnosus,* did not significantly induce Peyer’s patch enlargement in monocolonized mice (**Fig. 2, A to C**). In contrast, colonization with the commensal *E. coli* strain Nissle 1917 (EcN), a probiotic with a long history of use in humans (*25*, *26*), induced a significant increase in the Peyer’s patch area in monocolonized ex-germ-free mice (**Fig. 2, B and C**). Collectively, our results suggest that *E. coli* colonization triggers mucosal immune responses that promote Peyer’s patch enlargement and cellular recruitment to these lymphoid structures.

**Fig. 2.**
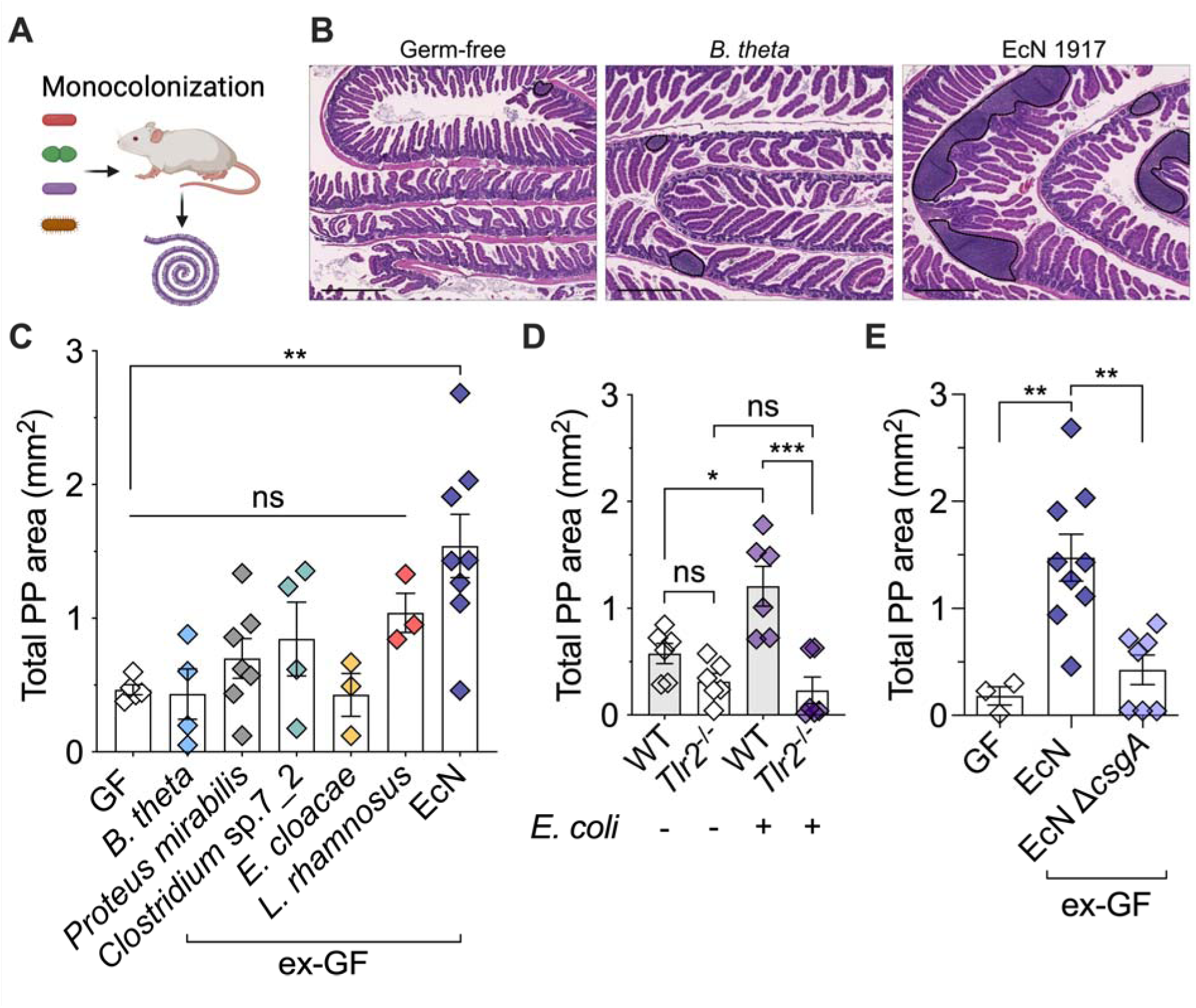
*E. coli* induces Peyer’s patch enlargement in germ-free mice through the expression of curli. **(A)** Experiment design. Germ-free (GF) mice were monocolonized with individual gut commensal strains for 7 days. Small intestinal tissues were processed as Swiss rolls for quantitative analysis of PP areas. **(B)** Representative H&E-stained sections from uncolonized GF mice and ex-GF mice colonized with *Bacteroides thetaiotaomicron* (*B. theta*) or *E. coli* Nissle 1917 (EcN). Black dashed lines delineate PP regions. 100X magnification, scale bars 500µm. **(C)** Quantitative analysis of PP areas in ex-GF mice monocolonized with the indicated commensal strains. **(D)** Quantification of PP areas in wild-type (*Tlr2*^+/+^) and *Tlr2*^-/-^ mice 7 days post-colonization with or without *E. coli* RRG1. **(E)** GF mice were either colonized with EcN or its curli-deficient mutant EcN Δ*csgA* for 7 days. PP areas were subsequently analyzed. Each symbol represents an individual mouse. Bars represent the mean ± SEM. Significant differences are indicated by *p ≤0.05, **p<0.01, ***p<0.001; ns = not significant. One-way ANOVA followed by Tukey’s multiple-comparison test. Illustrations were created using BioRender.

### Mice that lack Toll-like receptor 2 do not develop Peyer’s patches in response to *E. coli* colonization

We next sought to elucidate the mechanisms by which *E. coli* promotes the development of Peyer’s patches. In the gut, microbial components are primarily sensed by the intestinal epithelium through pattern recognition receptors, such as TLRs, which display region-specific expression patterns along the gastrointestinal tract (*11*). In the small intestine, TLR2 is the most uniformly expressed TLR across the intestinal epithelium in both adult and neonatal mice (*11*, *27–29*). Its expression spans the villi, crypts, and the follicle-associated epithelium of Peyer’s patches. To investigate whether TLR2 contributes to Peyer’s patch development, we colonized *Tlr2*^-/-^ mice and co-housed WT mice (Jax) with *E. coli* RRG1 and measured Peyer’s patch areas on Swiss roll sections after 7 days. *E. coli*-free *Tlr2*^-/-^ and co-housed WT mice (Jax) had comparably small Peyer’s patch areas (**Fig. 2D**). In contrast, *E. coli* colonization of WT mice induced a significant enlargement of Peyer’s patches, but this response was completely absent in *Tlr2*^-/-^ mice colonized with *E. coli* (**Fig. 2D**). These results indicate that TLR2 recognition of *E. coli* is essential for Peyer’s patch development.

### Bacterial curli amyloid is required for Peyer’s patch development

Given the requirement of TLR2 in *E. coli*-driven Peyer’s patch development, we next asked which *E. coli*-derived factor engages this pathway. In the gut, *E. coli* forms biofilms that promote mucosal adherence and facilitate persistent colonization (*30*, *31*). Key structural components of these biofilms (up to 85%) are curli amyloid fibers (*32*), which are recognized by the TLR2/TLR1 heterodimer (*33–35*). To test whether curli amyloid contributed to Peyer’s patch development, we colonized germ-free mice with the *E. coli* Nissle Δ*csgA* mutant, which lacks the major subunit of the curli fiber CsgA. Strikingly, *E. coli* Nissle Δ*csgA* failed to induce Peyer’s patch enlargement, despite similar colonization levels as *E. coli* Nissle wild-type and exhibited the characteristic morphotype observed under curli-inducing conditions (**Fig. 2E** and **fig. S2A** and **S2B**). Thus, the production of bacterial curli amyloid fibers contributes to *E. coli*-mediated Peyer’s patch development in adult mice.

### Abnormal B cell expansion and development in *Cxcr2^-^*^/-^ mice

We next sought to elucidate the cellular mechanisms driving B cell recruitment and accumulation in Peyer’s patches. Gene expression analysis of Peyer’s patches from *E. coli*-monocolonized ex-germ-free mice revealed a marked upregulation of the CXC chemokine genes *Cxcl1* and *Cxcl2*, with no corresponding induction in the spleen (**Fig. 3A**). These chemokines are canonical ligands for CXCR2, a receptor best known for its role in neutrophil trafficking (*36*). Surprisingly, *Cxcr2* expression was also elevated in Peyer’s patches, tissues typically devoid of neutrophils, while remaining low in the spleen (**Fig. 3B**). This unexpected expression pattern prompted us to investigate a potential role for CXCR2 in B cell recruitment. To this end, we longitudinally tracked CXCR2 expression in *E. coli*-monocolonized germ-free mice and found that transcriptional upregulation was accompanied by increased surface expression of CXCR2 on B cells within Peyer’s patches. Notably, CXCR2 peaked transiently on day 4 post-colonization, returning to low expression levels thereafter (**Fig. 3B**). This suggests a temporally regulated involvement in early B cell accumulation. Although CXCR2 is best characterized in the context of neutrophil chemotaxis, early studies on *Cxcr2^-^*^/-^ mice reported expansion of both neutrophils and B cells in the bone marrow, spleen, and peripheral lymph nodes, suggesting a broader role for CXCR2 beyond neutrophil trafficking (*37*). Thus, we analyzed the B cell distribution in primary and secondary lymphoid organs in *Cxcr2^-^*^/-^ mice. Despite relatively decreased B cell frequencies in the bone marrow (data not shown), absolute B cell numbers in *Cxcr2^-^*^/-^ mice were comparable to those of WT littermates (**fig. S3A**). Consistent with the original report, *Cxcr2^-^*^/-^ mice exhibited splenomegaly, characterized by an accumulation of B cells and neutrophils in the spleen (**fig. S3B**). Importantly, we did not observe a developmental block in peripheral B cell populations, as the numbers of transitional (T1 and T2), mature, or marginal zone (MZ) B cells in the spleen were comparable between WT and *Cxcr2^-^*^/-^ mice (**fig. S3E**). In the peripheral blood, we observed increased B cell counts in *Cxcr2^-^*^/-^mice compared to WT mice (**fig. S3C**). In contrast, mesenteric lymph nodes, which drain the small intestine (including Peyer’s patches) and the colon (*38*), exhibited comparable numbers of B cells between genotypes (**fig. S3D**).

**Fig. 3.**
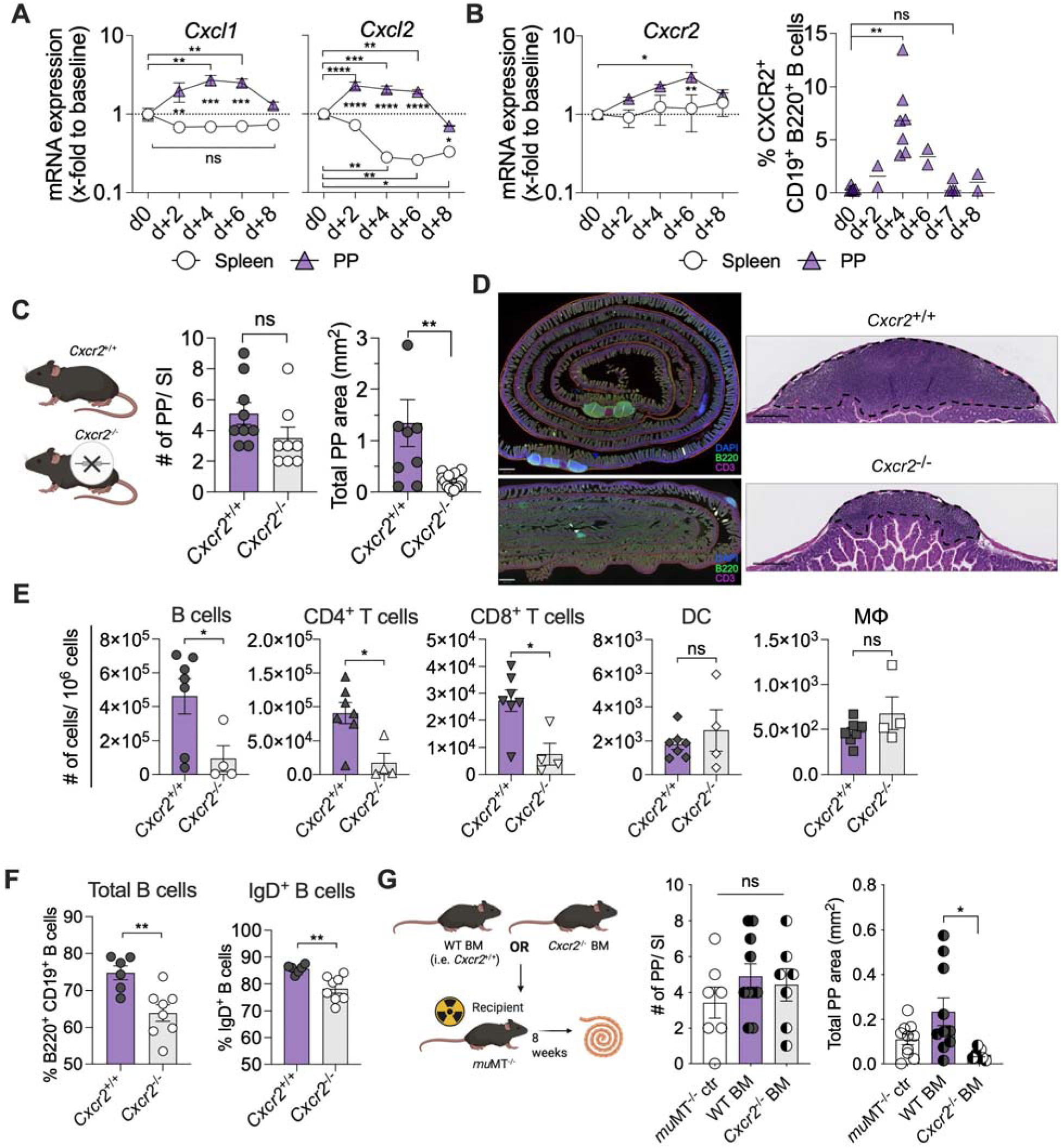
*Cxcr2^-/-^* mice exhibit hypoplastic Peyer’s patches with fewer naïve B cells. **(A)** GF mice were colonized with EcN, and the temporal expression of *Cxcl1* and *Cxcl2* mRNA was determined in whole tissue RNA from the spleen and Peyer’s patches (PP) over 8 days post-colonization. **(B)** Temporal analysis of *Cxcr2* mRNA expression in PP and spleen of EcN-colonized ex-GF mice, along with surface expression of CXCR2 on PP B cells. Statistical analysis for CXCR2 expression on PP B cells was performed only between timepoints with more than 3 datapoints (d0, d+4, d+7). (**C**) Quantitative analysis of PP numbers and areas of wild-type (*Cxcr2*^+/+^) and *Cxcr2*^-/-^ mice based on H&E-stained Swiss roll sections of the small intestine. (**D**) Representative small intestinal and immunofluorescently stained Swiss roll sections (20X magnification, scale bars 500µm) and H&E-stained PP (200X magnification, scale bars 250µm) from *Cxcr2*^+/+^ and *Cxcr2*^-/-^ mice. (**E**) Absolute numbers of immune cells isolated from PPs of *Cxcr2*^+/+^ and *Cxcr2*^-/-^ mice. B cells: B220^+^; CD4 T cells: CD3^+^CD4^+^; CD8 T cells: CD3^+^CD8^+^; DC: CD11b^+^ CD11c^+^ dendritic cells; MΦ: CD11b^+^ F4/80^+^ macrophages. (**F**) Frequencies of CD19^+^B220^+^ and IgD^+^ naïve B cells in PPs from *Cxcr2*^+/+^ and *Cxcr2*^-/-^ mice. (**G)** Experimental scheme. B-cell-deficient *mu*MT^-/-^ recipient mice were lethally irradiated and reconstituted with bone marrow cells from *Cxcr2*^+/+^ and *Cxcr2*^-/-^ donor mice. Following an 8-week recovery period, small intestinal tissues were processed as Swiss rolls for quantitative analysis of PP numbers and areas. Each symbol represents an individual mouse. Bars represent the mean ± SEM. Significant differences are indicated by *p ≤0.05, **p<0.01, ***p<0.001, ****p<0.0001, ns = not significant. Unpaired Student’s *t* test or one-way ANOVA followed by Tukey’s multiple-comparison test. Illustrations were created with BioRender.

### *Cxcr2^-^*^/-^ mice exhibit hypoplastic Peyer’s patches with fewer naive B cells

We then analyzed the small intestine of *Cxcr2^-^*^/-^ mice and their WT littermates. The overall number of Peyer’s patches per small intestine was similar between groups, indicating that CXCR2 does not appear necessary for embryonic Peyer’s patch organogenesis (*12*). However, Peyer’s patches in *Cxcr2^-^*^/-^ mice were hypoplastic and often difficult to detect macroscopically due to their reduced size and prominence (**Fig. 3, C and D**). Analysis of the cellular composition by flow cytometry identified a profound reduction of absolute B cell numbers as well as CD4^+^ and CD8^+^ T cells in *Cxcr2^-^*^/-^ mice, whereas other immune cell subsets (CD11b^+^ CD11c^+^ dendritic cells and CD11b^+^ F4/80^+^ macrophages) were not significantly different (**Fig. 3E)**. The relative frequency of Peyer’s patch B cells in *Cxcr2^-^*^/-^ mice was reduced by approximately 10%, which was largely attributed to a reduced frequency of IgD^+^ naive B cells compared to WT littermates (**Fig. 3F**), while the frequency of IgA^+^ cells was slightly increased (**fig. S3F**). The reduction in the number of CD4^+^ T cells was due to decreased frequencies of follicular T helper cells (TFH) but not follicular regulatory T cells (TFR; **fig. S3G**). *Cxcr2^-^*^/-^ mice also exhibited reduced frequencies and numbers of B cells in the colonic lamina propria, the effector site of the GALT, whereas T cell numbers were comparable between groups (**fig. S3H**). Expression levels of CXCR4 and 5, and α4β7, key molecules involved in B cell trafficking and organization within Peyer’s patches (*12*), were comparable between B cells from WT and *Cxcr2^-^*^/-^ mice (**fig. S3I**).

### CXCR2 contributes to B cell reconstitution in *mu*MT**^-/-^**mice

In a second approach, we performed bone marrow chimera studies to investigate the possible role of CXCR2 in B cell accumulation in Peyer’s patches. Recipient *mu*MT^-/-^ mice, which lack mature B cells (*39*), were lethally irradiated and reconstituted with bone marrow from *Cxcr2^-^*^/-^ or WT mice. After a reconstitution period of 8 weeks, we analyzed Peyer’s patches in Swiss roll sections from the small intestine of chimeric mice (**Fig. 3G**). The number of Peyer’s patch pockets was similar between groups. However, only WT bone marrow was sufficient to repopulate the Peyer’s patches in B cell-deficient recipients, whereas *Cxcr2^-^*^/-^ bone marrow failed to induce normal Peyer’s patch formation (**Fig. 3G**). Of note, recipient mice were not colonized with *E. coli,* which may account for the relatively moderate enlargement of Peyer’s patch areas observed in mice reconstituted with WT cells.

### CXCR2 blockade results in a reduced frequency of naive B cells in the gut of both adult and neonate mice

To control for possible developmental issues of *Cxcr2^-^*^/-^ mice, we next treated WT mice with rabbit anti-mouse CXCR2 serum (*40*, *41*) or control IgG and investigated the impact of CXCR2 blockade on the B cell pool in Peyer’s patches and the colonic lamina propria. Administration of CXCR2 antiserum to Taconic WT mice phenocopied observations from *Cxcr2^-^*^/-^ knockout mice, by decreasing the frequency and the total number of B cells in Peyer’s patches and the colonic lamina propria (**Fig. 4, A and B**). In Peyer’s patches, the depletion primarily affected IgD^+^ naive B cells. In contrast, T cell populations remained comparable between treatment groups (**Fig. 4, A and B**). Notably, B cell numbers in the spleen or mesenteric lymph nodes remained unaltered **(fig. S4A**), indicating that CXCR2 blockade was effective primarily on gut B cells. To investigate whether CXCR2 promotes Peyer’s patch development in response to *E. coli*, we monocolonized germ-free mice with *E. coli* Nissle and treated groups of mice with CXCR2 antiserum or control IgG (**Fig. 4C**). Seven days after colonization, we observed enlarged Peyer’s patches in mice colonized with *E. coli* Nissle that were treated with IgG control serum (**Fig. 4C**). Mice treated with CXCR2 antiserum showed a trend towards a reduction in Peyer’s patch size, although it did not reach statistical significance (**Fig. 4C**).

**Fig. 4.**
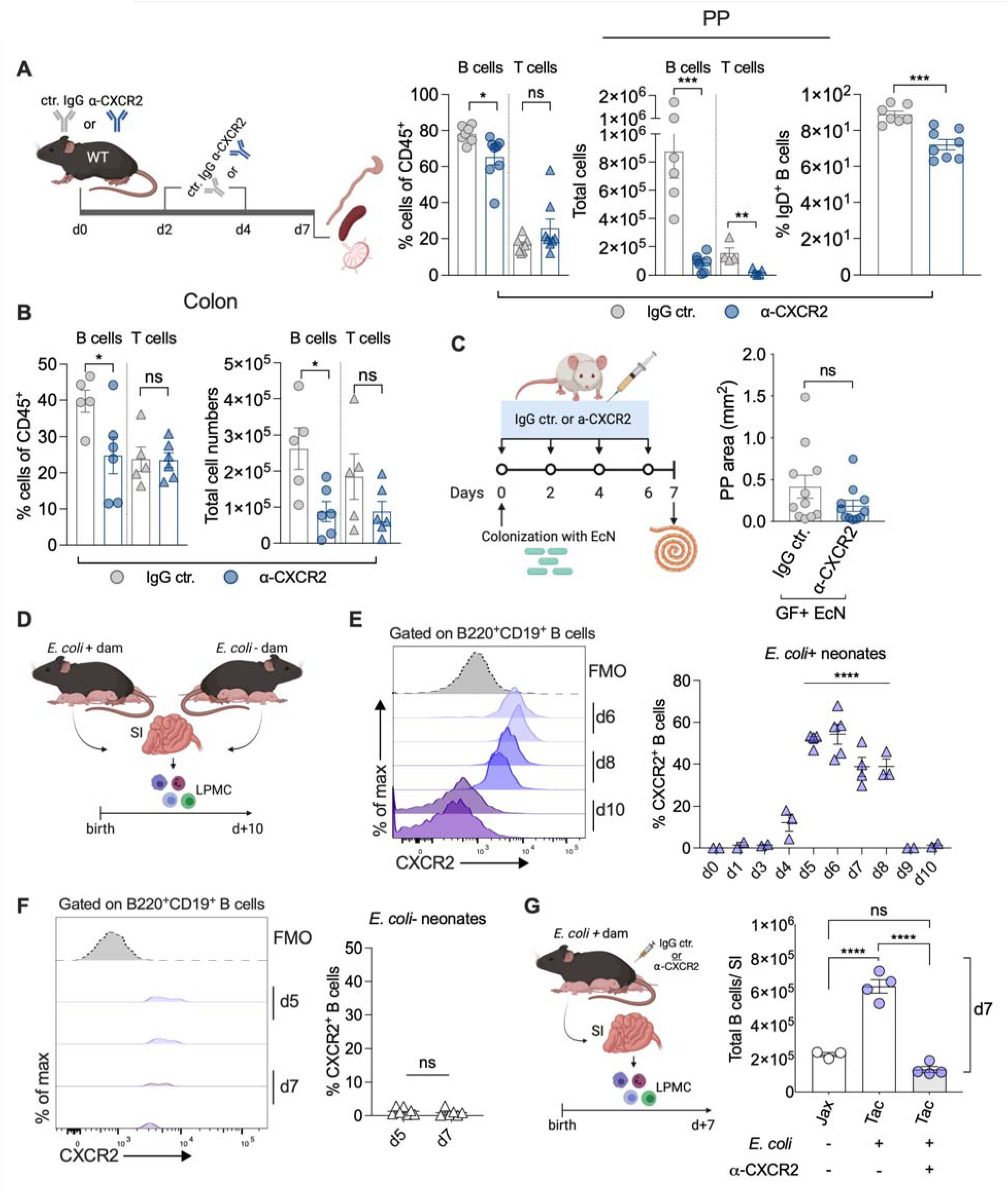
CXCR2 blockade leads to a decreased frequency of naive B cells in the intestinal mucosa. (**A**) Taconic mice were administered anti-CXCR2 serum or isotype control serum (IgG) intraperitoneally (i.p.) on days 0, 2, and 4. CD19^+^B220^+^ B cell and CD3^+^ T cell populations in (**A**) Peyer’s patches (PP) and (**B**) the colonic lamina propria were subsequently quantified by flow cytometry. (**C**) Germ-free mice were colonized with *E. coli* Nissle 1917 (EcN) and given anti- CXCR2 serum or rabbit IgG control serum starting on day 0, and administered i.p. every 48 hours until day 6. Small intestines were processed as Swiss rolls on day 7 and analyzed for PP area. (**D**, **E**, and **F**) CXCR2 expression on small intestinal B cells was longitudinally assessed in the offspring of Taconic (**E**: *E. coli* +) and Jackson (**F:** *E. coli* -) dams. d# = Days post-birth. (**G**) Taconic dams received i.p. injections of either anti-CXCR2 or isotype control (IgG) serum immediately after parturition, and administration was repeated every 48 hours until day 6. On postnatal day 7, lamina propria mononuclear cells (LPMCs) were isolated from the small intestines of the offspring, and B cells were quantified by flow cytometry. Results were compared to age-matched pups from untreated Jackson dams. Bars represent the mean ± SEM. Significant differences are indicated by *p ≤0.05, ***p<0.001, ****p<0.0001, ns = not significant. Unpaired Student’s *t* test or one-way ANOVA followed by Tukey’s multiple-comparison test. Illustrations were created using BioRender.

### B cell accumulation in the neonatal gut is dependent on *E. coli* colonization and CXCR2

Next, we investigated whether *E. coli*-dependent accumulation of B cells during the early postnatal period (**Fig. 1G**) requires CXCR2 signaling. To this end, we first assessed CXCR2 expression on small intestinal lamina propria B cells in offspring of *E. coli*-positive Taconic or *E. coli*-negative Jackson mice over time (**Fig. 4D**). In pups born to Taconic dams, CXCR2^+^ B cells were detectable, with expression peaking around postnatal day six (**Fig. 4E**). In contrast, CXCR2 expression was absent on lamina propria B cells from Jackson pups lacking *E. coli* (**Fig. 4F**). To functionally test the role of CXCR in B cell accumulation in the neonatal gut, we administered CXCR2 antiserum to Taconic dams immediately after parturition, anticipating passive transfer of circulating maternal IgG to the pups via the dam’s milk (*42*). At day 7, pups from *E. coli*-positive dams treated with control IgG had significantly more lamina propria B cells than *E. coli*-negative pups from Jackson dams (**Fig. 4G**). Remarkably, pups fostered by *E. coli*+ Tac dams receiving CXCR2 antiserum treatment had lower gut B cell numbers, comparable to pups from *E. coli*-negative dams (**Fig. 4G**). Collectively, these data indicate that *E. coli* colonization triggers intestinal B cell accumulation in neonates in a CXCR2-mediated manner.

### Targeted deletion of *Cxcr2* in B cells leads to diminished Peyer’s patch formation and reduced B cell accumulation following *E. coli* colonization

To investigate the intrinsic role of CXCR2 in B cells, we crossed *Cxcr2*^fl/fl^ mice (*43*) with hCD20-Cre (Tam-hCD20-Cre) mice (*44*), a tamoxifen-inducible Cre model that enables B cell-specific Cre activation. This approach allows the evaluation of the impact of selective CXCR2 depletion in B cells following tamoxifen administration and *E. coli* colonization. We first validated Cre expression in B cells across various organs, including the bone marrow, spleen, and Peyer’s patches, by crossing the hCD20Tam-Cre mouse with the ROSA26^tdTomato^ reporter strain (*45*). While Cre induction in bone marrow B cells was more heterogeneous (∼30%), Cre activity in splenic and Peyer’s patch B cells remained consistent across samples (∼67% and ∼63% respectively; **fig. S5A**). Cre expression in non-B cells was negligible (**fig. S5A**). *Cxcr2*^fl/fl^ x hCD20-Cre^+^ and *Cxcr2*^fl/fl^ x hCD20-Cre^-^ littermates received intraperitoneal tamoxifen injections prior to colonization with *E. coli* RRG1 (**Fig. 5A**). Colonization levels were comparable between the groups throughout the experiment (**fig. S5B**). *Cxcr2*^fl/fl^ x hCD20-Cre^+^ exhibited significantly smaller Peyer’s patch areas compared to their *Cxcr2*^fl/fl^ x hCD20-Cre^-^ littermate controls (**Fig. 5B**). This reduction in overall area was accompanied by a decrease in both B cell frequencies and absolute B cell numbers (**Fig. 5, C and D**). Additionally, we observed a trend toward reduced IgD+ B cell frequencies, resulting in a significant decrease in the absolute number of naive IgD+ B cells in Peyer’s patches of *Cxcr2*^fl/fl^ x hCD20-Cre^+^ mice (**Fig. 5E**). Similarly, while the frequencies of IgA+ B cells did not differ significantly between groups, their absolute numbers were reduced (**fig. S5C**). In contrast, despite increased T cell frequencies in *Cxcr2*^fl/fl^ x hCD20-Cre^+^ mice, the absolute T cell numbers remained comparable between both genotypes, showing no significant differences (**Fig. 5C and fig. S5D**). We conclude that targeted deletion of CXCR2 in B cells reduces B cell accumulation in Peyer’s patches in response to *E. coli* colonization.

**Fig. 5.**
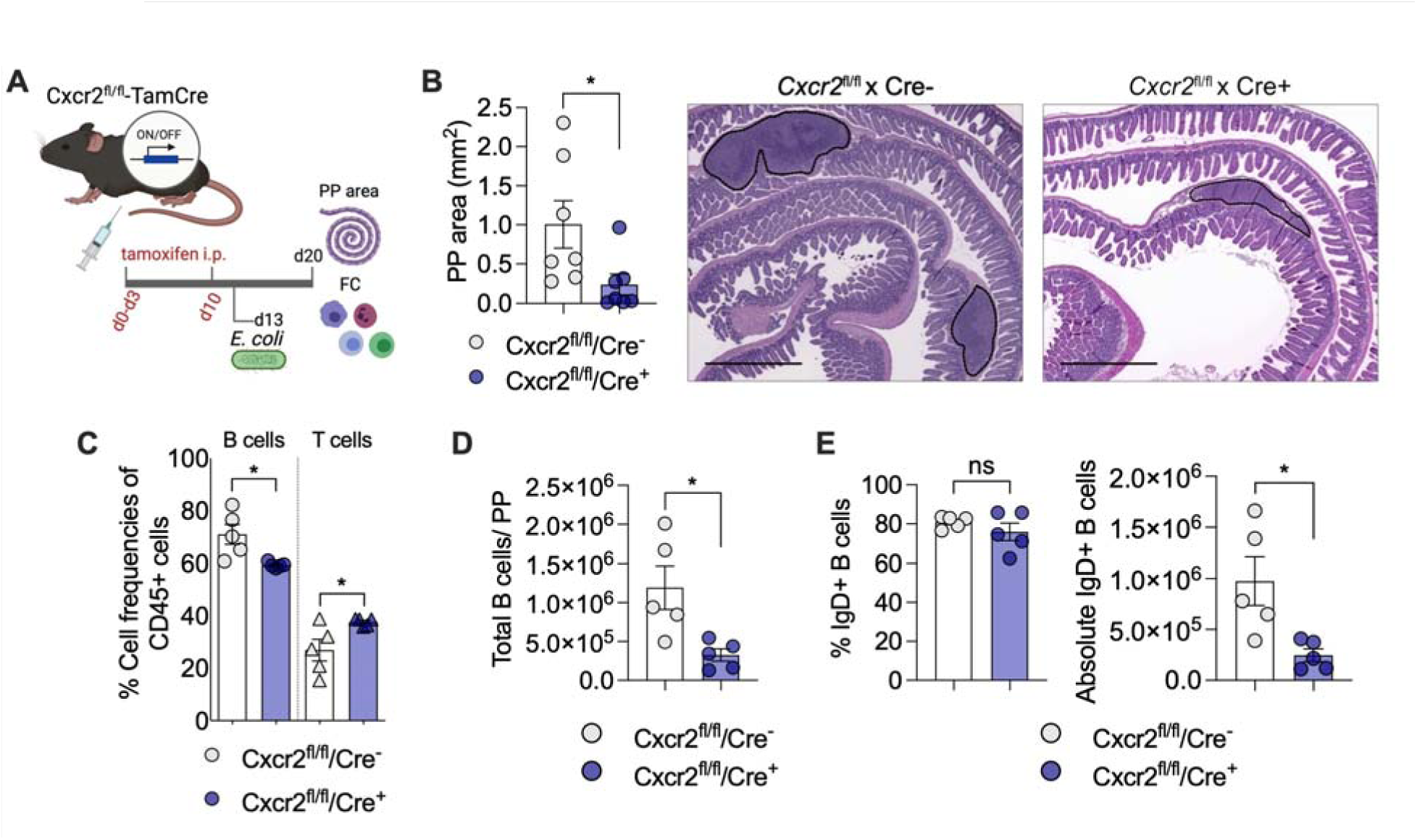
Targeted deletion of *Cxcr2* in B cells leads to diminished Peyer’s patch formation with reduced B cell accumulation after *E. coli* colonization. (A) *Cxcr2*^fl/fl^ x hCD20-TamCre^+^ and *Cxcr2*^fl/fl^ x hCD20-TamCre^-^ mice received intraperitoneal (i.p.) injections of tamoxifen as indicated in the scheme (d = day) to induce B cell-specific *Cxcr2* deletion. On day 13, mice were colonized with *E. coli* RRG1. Seven days post-colonization, small intestines were harvested and processed as Swiss rolls for histological analysis. (**B**) Quantification of Peyer’s patch (PP) areas of *Cxcr2*^fl/fl^/Cre^+^ and *Cxcr2*^fl/fl^/Cre^-^ mice, along with representative H&E-stained sections. 50X magnification, scale bars 1mm. (**C**) Flow cytometric analysis of the frequencies of CD19^+^B220^+^ B cells and CD3^+^ T cells and (**D**) absolute numbers of B cells in PPs from *Cxcr2*^fl/fl^/Cre^+^ and *Cxcr2*^fl/fl^/Cre^-^ mice following tamoxifen-induced gene deletion and *E. coli* colonization, as outlined in (A). (**E**) Frequencies and absolute numbers of IgD^+^ naïve B cells in PP from *Cxcr2*^fl/fl^/Cre^+^ and *Cxcr2*^fl/fl^/Cre^-^ mice as determined by flow cytometry. Bars represent the mean ± SEM. Significant differences are indicated by *p ≤0.05, ns = not significant. Unpaired Student’s *t* test. Illustrations were created using BioRender.

## Discussion

Microbial stimulation is a key driver of immune cell accumulation in the gut, and the intestinal microbiota plays a fundamental role in shaping the postnatal development of the mucosal immune system. However, the specific microbial factors that influence the magnitude, composition, and spatial organization of mucosal immune responses remain poorly defined.

In this study, we identify *E. coli* as a potent inducer of B cell accumulation within Peyer’s patches and the lamina propria of the small intestine. This process is mediated by the expression of curli, biofilm-associated amyloid fibers, and requires host sensing via TLR2. Neither curli-deficient *E. coli* strains nor TLR2-deficient hosts exhibited B cell recruitment or the associated Peyer’s patch expansion, indicating a critical role for this microbe-host interaction in shaping local immune architecture. We further show that B cell accumulation in response to *E. coli* colonization is dependent on the chemokine receptor CXCR2, which is traditionally associated with neutrophil trafficking. Notably, the human CXCR2 homolog - interleukin-8 receptor (IL-8R) - has been reported to be expressed on human B cells, with increased expression levels observed during HIV infection (*46*). Furthermore, IL8R+ B cells are capable of migrating along IL-8 chemokine gradients (*46*, *47*), suggesting a conserved role for this chemokine axis in directing B cell positioning during inflammation or microbial challenge. Consistent with this, colonization of germ-free mice with *E. coli* Nissle 1917 led to the selective upregulation of the chemokines *Cxcl1* and *Cxcl2* within Peyer’s patches, but not in the spleen, suggesting that localized chemokine induction may contribute to B cell trafficking at mucosal sites. In contrast to such inflammation-induced recruitment, homeostatic lymphocyte trafficking to, within, and from Peyer’s patches is orchestrated by tightly regulated and sequential interactions between integrin-adhesion molecule pairs and chemokine receptor-ligand axes. Key examples include α4β7-MAdCAM-1, CXCR4-CXCL12, CXCR5-CXCL13, and CCR7-CCL21, which coordinate the migration and positioning of B and T cells through high endothelial venules and stromal compartments within Peyer’s patches (*12*).

Our findings may have broader relevance for immunological and biomedical research, as we found that the presence or absence of *E. coli* had a profound influence on mucosal immune architecture in mice. This highlights the importance of microbial composition as a variable in experimental design. A notable example is a previous study comparing B6 mice from Jackson Laboratories and Taconic Farms, which revealed that Th17 cell differentiation was induced by the presence of segmented filamentous bacteria (SFB) (*4*), a commensal microbe absent in Jackson mice. *E. coli*, a common pioneer colonizer of the neonatal gut in most warm-blooded animals, has been primarily studied in the context of niche occupation and competition with enteric pathogens such as *Salmonella* (*19*, *48*, *49*), but our findings suggest an additional immunomodulatory role. From an evolutionary standpoint, the recruitment of B cells in response to early-life *E. coli* colonization may serve as a host defense mechanism to contain the microbe’s expansion and prevent systemic dissemination, thereby reinforcing mucosal immune barriers against this potentially opportunistic species. Supporting this notion, a study in germ-free recombinase-activating gene 1 (*Rag^-^*^/-^) mice (lacking both T and B cells) showed that monocolonization with ‘probiotic’ *E. coli* Nissle 1917 led to 100% mortality, whereas colonization under SPF conditions was well tolerated (*50*). Indeed, commensal *E. coli* is harmless and even beneficial in healthy individuals, but can cause potentially life-threatening infections like sepsis or neonatal meningitis in vulnerable populations, including cancer patients, the elderly, or preterm infants (*51–53*). Likewise, curli expression is beneficial in the gut, where it plays immunomodulatory roles and promotes barrier function (*54*), but can also be detrimental in specific contexts. A clinical study in human sepsis patients identified anti-CsgA (curli) antibodies and higher circulating proinflammatory cytokines in convalescent individuals but not in healthy controls, suggesting that curli expression may elicit systemic immune responses during invasive infection (*55*).

Our findings provide important insights into how microbial components shape mucosal immunity and reveal a direct link between *E. coli* colonization and B cell recruitment to the gut. Future studies aimed at elucidating the functional consequences of *E. coli-*driven Peyer’s patch maturation will further advance our understanding of the complex interplay between commensal microbes and the mucosal immune system, with broad implications for health and disease.

## Acknowledgements

Confocal imaging and slide scanning were done at the UCSD School of Medicine Microscopy Core, which is supported by a NINDS P30 grant (NS047101). Flow cytometry experiments were performed either at the La Jolla Institute for Immunology or at the Sanford Burnham Prebys Flow Cytometry Core. Histology was performed at the La Jolla Institute for Immunology core and was partly supported by NIDDK Grant P30 DK120515.

## Funding

Max Kade Foundation (RRG)

Crohn’s and Colitis Foundation grant 649744 (RRG)

National Institute of Health grant T32AI007036 (GTW)

American Heart Association (SMK)

National Institute of Health grant R01AI114625 (MR)

National Institute of Health grant R37AI126277 (MR)

National Institute of Health grant R01AI108651 (L-FL)

Burroughs Wellcome Fund Investigator in the Pathogenesis of Infectious Disease Award (MR)

Kenneth Rainin Foundation (MR and HC)

AMED grant JP233fa627004 (MR and HC)

Chiba University-University of California-San Diego (UCSD)

Center for Mucosal Immunology, Allergy, and Vaccines (MR and HC)

## Author contributions

Conceptualization: RRG, MR

Methodology: RRG, GTW, SMK, KM, RMR, DDL, TEL, AR, S-PN, JB, EIZ, L-FL, ÇT, HC, MR

Investigation: RRG, GTW, SMK, KM, SH, KS, C-YH, FJA, WS, RP, MC-T, GJN, FT, AP-L, PS, MPW, VL, DDL, JB

Resources: RMR, TEL, AR, EIZ, L-FL, ÇT, HC

Visualization: RRG Formal analysis: RRG

Funding acquisition: MR, HC, JB, LL Project administration: MR

Supervision: RRG, S-PN, JB, EIZ, L-FL, ÇT, HC, MR

Writing – original draft: RRG, MR

Writing – review & editing: RRG, GTW, RP, GJN, FT, AP-L, MPW, RMR, S-PN, JB, EIZ, L-FL, CT, HC, MR

## Data and materials availability

All data are available in the main text or in the supplementary materials

## Supplementary Materials

Materials and Methods

Figs. S1 to S5

Tables S1 to S2

References (1–15)

