## Supplemental files for "Commensal *Escherichia coli* colonization triggers Peyer’s patch development"

### SUPPLEMENTAL MATERIALS

### Materials and Methods

#### Mice

C57BL/6 wild-type (WT), *Ighm*$^{tm1Cgn}$ (*mu*MT$^{-/-}$), and *Tlr2*$^{-/-}$ mice, as well as the ROSA26$^{tdTomato}$ reporter strain mice, were obtained from The Jackson Laboratory. Additional C57BL/6 WT mice were purchased from Taconic Farms. *Cxcr2*$^{-/-}$ mice (*1*) were initially provided by T.E. Lane (UC Irvine, C57BL/6 background) and subsequently crossed with C57BL/6 Nramp1$^{+}$ (*2*) from G. Barton (UC Berkeley). The crossing was necessary because the *Cxcr2* gene is in close proximity to the *Slc11a1* gene encoding Nramp1, and the embryonic stem cells used to make the *Cxcr2* mutant were derived from 129 mice, which are Nramp1+ (*3*). *Cxcr2*$^{+/+}$ Nramp1$^{+}$ and *Cxcr2*$^{-/-}$ Nramp1$^{+}$ were generated by intercrossing littermates of heterozygous parents. *Cxcr2*$^{fl/fl}$ mice were initially described by Liu et al (*4*). To enable B cell-specific deletion, *Cxcr2*$^{fl/fl}$ were crossed with hCD20-TamCre (MS4A1-cre/ERT2) mice (*5*), kindly provided by M. Shlomchik (University of Pittsburgh). This strain expresses a tamoxifen-inducible Cre recombinase under control of the MS4A1 promoter, enabling conditional gene deletion in B cells. Tamoxifen-induced conditional deletion of *Cxcr2* was achieved by administering daily intraperitoneal injections of 3 mg tamoxifen (dissolved in pre-heated, sterile-filtered corn oil) to 5-6-week-old mice for four consecutive days. This was followed by a 7-day rest period. To enhance recombination efficiency, mice then received a single tamoxifen booster dose, followed by a 3-day rest period, then colonization with RRG1 (see scheme in **Fig. 5A**). In experiments involving ROSA26$^{tdTomato}$ reporter mice, organs were harvested 7 days after the final tamoxifen injection. All mice were bred and maintained under specific pathogen-free (SPF) conditions in the vivarium at the University of California, San Diego. Germ-free C57BL/6 mice were bred in specific isolators at the University of California, San Diego, and their germ-free status was confirmed by aerobic and anaerobic culture, Gram staining of feces, and quantitative real-time PCR (qRT-PCR) of 16S rDNA. All experimental procedures were approved by the Institutional Animal Care and Use Committee (IACUC) at UC San Diego or UC Irvine.

#### Colonization experiments

For colonization experiments, *E. coli*-negative adult mice (male and female, 6-8 weeks of age) were orally gavaged with 20mg streptomycin/mouse, followed by $10^9$ CFU of *E. coli* RRG1 or *E. coli* Nissle 1917 24 hours later, or provided with respective *E. coli* strains in the drinking water for 2 days (approximately $10^7$ CFU/mL). Intestinal colonization was confirmed by plating fecal samples on MacConkey agar. Germ-free mice (6-8 weeks of age) were monocolonized via oral gavage with $10^9$ CFU of the bacterial strains listed in Table S1 and maintained for 7 days in isocages. At the end of the experiments, organs were harvested and Swiss rolls prepared as described below.

**CXCR2 antiserum treatment in adult and neonate mice**

The CXCR2 antiserum was developed and validated as previously described (*6*). For depletion experiments in germ-free mice, 200 µL of filter-sterilized CXCR2 antiserum was administered intraperitoneally starting on the day of colonization with $10^8$ CFU of *E. coli* Nissle, and repeated every 48 hours until day 6 (*i.e.*, a total of 4 injections). Organs were harvested on day 7 of the experiment. Control animals received equivalent volumes of polyclonal rabbit IgG (Jackson ImmunoResearch). In other experiments, 8-week-old Taconic mice were given 200 µL of filter-sterilized CXCR2 antiserum or polyclonal rabbit IgG every other day for a total of three times, and organs were harvested on day 7 of the experiment. For neonatal depletion experiments, Taconic dams were administered 300 µL of CXCR2 antiserum every other day beginning after parturition and continuing every 48 hours until day 6. Lamina propria mononuclear cells (LPMCs) from the small intestine of neonates were isolated on day 7 of the experiments.

**Bone marrow chimera experiments**

6-8 week-old *mu*MT$^{-/-}$ recipient mice were irradiated with 10 Gray using an X-ray energy irradiator. The following day, bone marrow (BM) cells were harvested from *Cxcr2*$^{+/+}$ or *Cxcr2*$^{-/-}$ donor mice by flushing femurs with sterile, ice-cold PBS. Red blood cells were lysed with ACK lysis buffer, and BM cells were counted and adjusted to $5 \times 10^7$ cells/ mL PBS. Irradiated recipients were then reconstituted with $5 \times 10^6$ BM cells/ mouse intravenously. Peyer's patch formation was evaluated 8 weeks after BM transplantation on Swiss roll sections as described below.

**Swiss roll preparation, histological analysis, and immunofluorescence**

Swiss Rolls were prepared from the small intestine. The intestine was divided into three equal segments, and luminal contents were flushed out using 10% neutral-buffered formalin. Each segment was opened longitudinally and rolled from the proximal end with the luminal side facing inward, using a toothpick. Swiss rolls and other tissue samples were fixed in 10% neutral-buffered formalin for 24-48 hr, then transferred to 70% ethanol until further processing. Tissues were embedded in paraffin according to standard protocols. Serial sections (5 µm sections) were cut until the entire Swiss roll was represented and stained with hematoxylin and eosin (H&E). Slides were scanned using either a Nanozoomer 2.0 HT (Hamamatsu) or a Zeiss Axio Scan.Z1 slide scanner. Peyer's patches were identified at 100 x magnification and annotated using the freehand region tool in NDP.view2 (Hamamatsu) or QuPath software. The area of each Peyer's patch was measured in mm$^2$, and total Peyer's patch area per animal was calculated by summing all annotated regions. For immunofluorescence stainings, deparaffinized Swiss roll sections were subjected to antigen retrieval using sodium citrate buffer (pH 6.0) for 30 minutes (min) in a pre-heated food steamer, followed by a 20-min cooling period at room temperature. Endogenous peroxidase activity was quenched using Dako Dual Endogenous Enzyme Block (Agilent) for 10 min. After washing with TBS containing 0.1% Tween-20 (T-BST), non-specific binding was blocked with Agilent Protein Block. Sections were then incubated overnight at 4°C in a humidified chamber with primary antibodies against B220 and CD3ε (see Table S2). The following day, slides were washed and incubated with fluorophore-conjugated secondary antibodies (see Table S2) for 2 h at room temperature. After final washes, slides were coverslipped with ProLong Gold Antifade Mountant containing DAPI (Thermo Fisher Scientific), allowed to dry, and imaged using an Olympus Slideview VS200 system.

#### Immune cell isolation from Peyer's patches, mesenteric lymph nodes, and intestinal lamina propria

For isolation of immune cells from mesenteric lymph nodes (MLN) or Peyer's patches (PP), mice were euthanized, and the small intestine was carefully removed. MLNs were dissected, and PP were visually identified along the antimesenteric side of the intestine. PP were excised using curved surgical scissors, avoiding surrounding tissue, and immediately transferred into ice-cold PBS. To generate single-cell suspensions, tissues were placed onto 70 µm nylon mesh cell strainers and mechanically dissociated by gently pressing through the mesh using the plunger end of a 1 mL syringe. Cells were washed through the strainer with PBS and collected in 50 mL conical tubes. Viable cells were counted using a Countess II automated cell counter (Invitrogen) prior to downstream applications. To isolate lamina propria mononuclear cells (LPMCs) from the colon or the small intestine of neonates, the intestine was harvested, flushed with ice-cold PBS, and opened longitudinally. The tissue was minced using a scalpel and incubated in calcium- and magnesium-free Hank's balanced salt solution (HBSS) supplemented with 10% fetal bovine serum (FBS), 1mM dithiothreitol (DTT), and 2mM EDTA for 20 min at room temperature under continuous shaking. Subsequently, the fragments were vortexed at full speed for 4 min, the supernatants discarded, and the remaining fragments were resuspended in HBSS/FBS. A second vortexing step was performed for 2 min to further remove epithelial cells, followed by supernatant removal. The remaining tissue was transferred into Iscove's Modified Dulbecco's Medium (IMDM; Life Technologies) supplemented with 20% FBS, 128 U/mL type VIII collagenase, and 10 U/mL DNase I (both Sigma-Aldrich), and digested for 60 min at 37°C with gentle rotation. Following enzymatic digestion, the samples were thoroughly resuspended and sequentially filtered through 100µm, 70µm, then 40 µm nylon mesh cell strainers. Cell suspensions were centrifuged at 350 × *g* for 10 min at 4°C, and viable cells were counted using an automated cell counter before use in downstream applications.

#### Immune cell isolation from the spleen and blood

Splenocytes were isolated by excising the spleen and mincing the tissue into small pieces in 10 mL RPMI medium using a scalpel in a sterile petri dish. The tissue fragments were then transferred onto a 70 µm nylon mesh cell strainer and mechanically disrupted using the plunger end of a 1 mL syringe, followed by flushing with an additional 10 mL of RPMI. The resulting single-cell suspension was centrifuged at 350 × *g* for 5 minutes at 4°C. After discarding the supernatant, the cell pellet was resuspended in 4 mL of ice-cold 1x red blood cell (RBC) lysis buffer and incubated for 5 minutes on ice. Cells were then washed with 20 mL of cold PBS, centrifuged again, and the pellet resuspended in 10 mL PBS. Viable cells were counted using trypan blue exclusion prior to downstream applications. Bone marrow cells were isolated by cutting open the ends of the femurs with sterile scissors, followed by flushing the marrow cavity with ice-cold PBS. The resulting cell suspension was thoroughly resuspended, filtered through a 70 µm cell strainer, and centrifuged at 350 × g for 5 minutes at 4°C. RBC lysis was performed as described above, and viable cells were counted using an automated cell counter. For blood cell isolation, whole blood was collected via cardiac puncture of the right atrium using a pre-heparinized syringe. A volume of 300µL was transferred into a 15 mL conical tube containing 5 mL RBC lysis buffer and incubated on ice for 5 minutes. The lysis reaction was quenched with PBS, followed by centrifugation at 350 × g for 5 minutes at 4°C. The supernatant was discarded, and the cell pellet was resuspended in PBS for viable cell counting and downstream analyses.

#### Flow cytometry experiments

For flow cytometry, single-cell suspensions were prepared from tissue samples and resuspended in ice-cold PBS. Cells were stained with a Zombie Yellow viability dye (see Table S2) and incubated with an anti-CD16/CD32 (Fc block, see Table S2) for 7 minutes on ice to minimize non-specific Fc receptor binding. After two washes with flow buffer (PBS containing 1% bovine serum albumin), cells were labeled with fluorophore-conjugated surface antibodies for 30 min on ice (see Table S2). All samples were fixed in 4% Paraformaldehyde for 20 min at RT prior to acquisition. In some experiments, surface staining was followed by intracellular staining using the True-Nuclear Transcription Factor Buffer Set (BioLegend) according to the manufacturer's instructions. Data were acquired using an LSRFortessa flow cytometer (BD Biosciences) or a Sony SA 3800 and analyzed with FlowJo software (version 10).

**RNA Isolation, reverse transcription, and quantitative PCR**

Tissue samples were harvested and immediately stored in RNA*later* (Invitrogen) until further processing. For total RNA extraction, tissues were lysed in RLT buffer, and RNA was isolated using the RNeasy Mini Kit (Qiagen) according to the manufacturer's instructions. Reverse transcription was performed using the SuperScript IV VILO Master Mix (Invitrogen) to generate cDNA. Gene expression was assessed by quantitative real-time PCR using the PowerUp SYBR Green Master Mix on a QuantStudio 5 Real-Time PCR System (both Thermo Fisher Scientific). Data were analyzed using the comparative $2^{-\Delta\Delta Ct}$ method, with target gene expression normalized to *Actb* (β-actin) mRNA levels. Primer sequences used were as follows: *Actb* forward: 'GGCTGTATTCCCCTCCATCG3′ and reverse: 5′CCAGTTGGTAACAATGCCATGT3′; *Cxcr2* forward: 'ATGCCCTCTATTCTGCCAGAT' and reverse: 'GTGCTCCGGTTGTATAAGATGAC'; *Cxcl1* forward: 'TGCACCCAAACCGAAGTCAT' and reverse: 5'TTGTCAGAAGCCAGCGTTCAC'; *Cxcl2* forward: 'TGCCGGCTCCTCAGTGCTG ' and reverse: 'AAACTTTTTGACCGCCCTTGA'.

**Whole genome sequencing**

Genomic DNA was extracted from *E. coli* RRG1 broth culture using the Qiagen DNeasy Blood & Tissue Kit according to the manufacturer's instructions. Hybrid shotgun and Oxford Nanopore Technology sequencing, genome assembly, and annotation (Bakta v1.11) were performed by Plasmidsaurus. For phylogenetic comparison of *E. coli* RRG1 in the context of other *Escherichia spp.*, an assembly set consisting of *E. coli* RRG1, the top 2 *Escherichia coli* NCBI BLASTn hits to RRG1 (*7*), 100 *Escherichia coli* genomes randomly selected from 10,401 complete assemblies in the NCBI genome collection, 12 representative genomes for different *E. coli* phylogroups (*8*), and the *Escherichia albertii* RefSeq reference genome (GCF_028622335.1) was constructed (NCBI datasets v17.3.0) (*9*). Mash distances were calculated between the 116 genomes in the assembly set (Mash v2.3) (*10*), and a phylogenetic tree was constructed (R v4.4.2; ape v5.8.1). The phylogenetic tree was then rooted to *Escherichia albertii* and visualized (iTOL v7.2.1) (*11*), and phylogroup annotations were assigned based on phylogroup representatives included in the assembly set.

**Biofilm morphotype**

For visual characterization of curli expression, bacterial strains were grown at 28 °C for 72 h on YESCA agar plates supplemented with 40 µg/mL Congo Red and 20 µg/mL Coomassie Blue (*12*).

**Statistical analysis**

Statistical analyses were performed using GraphPad Prism v10. Colony-forming unit (CFU) data were log-transformed prior to statistical testing to normalize variance. The distribution of data was assessed using the Shapiro-Wilk test. Outliers were identified and removed with the ROUT test. For comparisons between two groups with normally distributed data, unpaired Student's *t*-tests were applied. For non-normally distributed unpaired datasets, the Mann-Whitney *U* test was used. When comparing more than two groups, one-way analysis of variance (ANOVA) followed by Tukey's multiple-comparisons test was conducted.

**Materials availability statement**.
Mice, bacterial strains, and reagents generated in this study will be provided by the corresponding author upon reasonable request and processing of a material transfer agreement.

### SUPPLEMENTAL FIGURES

#### Supplemental Figure 1

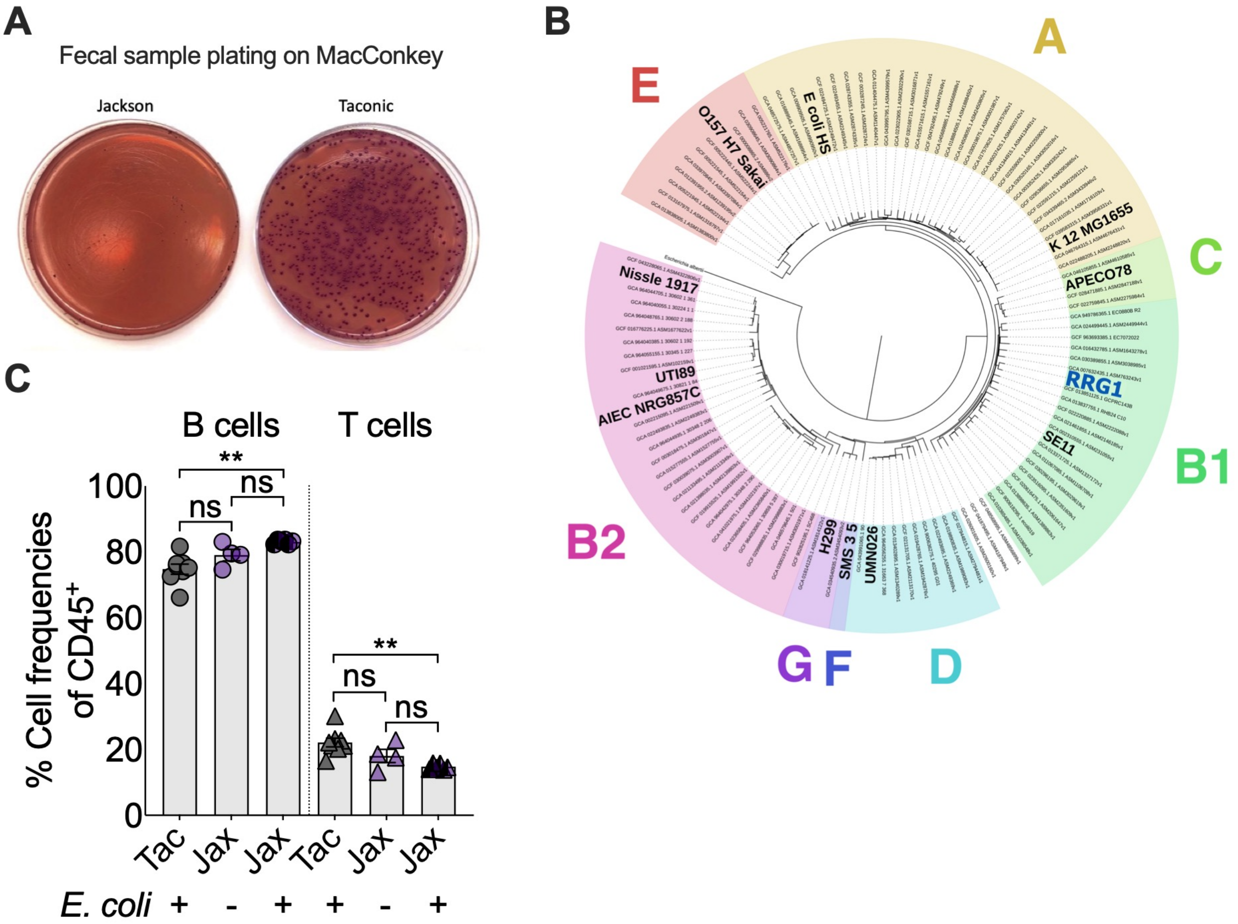


**fig. S1. Mice from Taconic are colonized with commensal *E. coli***
**(A)** Fecal pellets from Jax and Tac mice were homogenized in PBS and plated on MacConkey agar overnight. Lactose-positive colonies were observed on plates from Tac mice, identified as *E. coli,* and designated RRG1. **(B)** Phylogenetic tree of RRG1 whole genome sequence compared to n=115 publicly available *E. coli* or *E. albertii* (root) genomes using Mash (*7*). **(C)** Cell frequencies of $CD19^{+}B220^{+}$ B cells and $CD3^{+}$ T cells from Peyer's patches of Taconic and Jackson mice with and without *E. coli.* Significant differences are indicated by **p ≤0.01, ns = not significant. One-way ANOVA followed by Tukey's multiple-comparison test. Data represent the mean ± SEM.

### Supplemental Figure 2

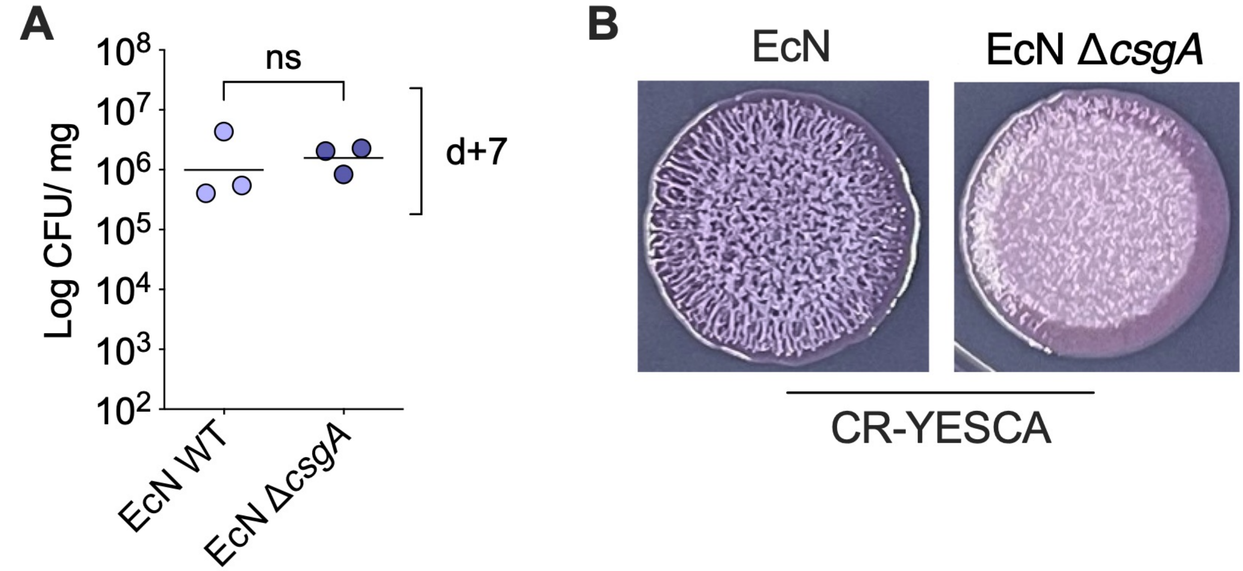


**fig. S2. Colonization of *E. coli* Nissle 1917 wild-type and the Δ*csgA* mutant in germ-free mice. (A)** Fecal colony-forming units (CFU) per mg feces on day 7 after monocolonization of germ-free adult mice. **(B)** Colony morphology of EcN wild-type and Δ*csgA* mutant grown on Congo Red (CR)-YESCA agar. Data represent the geometric mean; ns = not significant. Mann-Whitney *U* test.

### Supplemental Figure 3

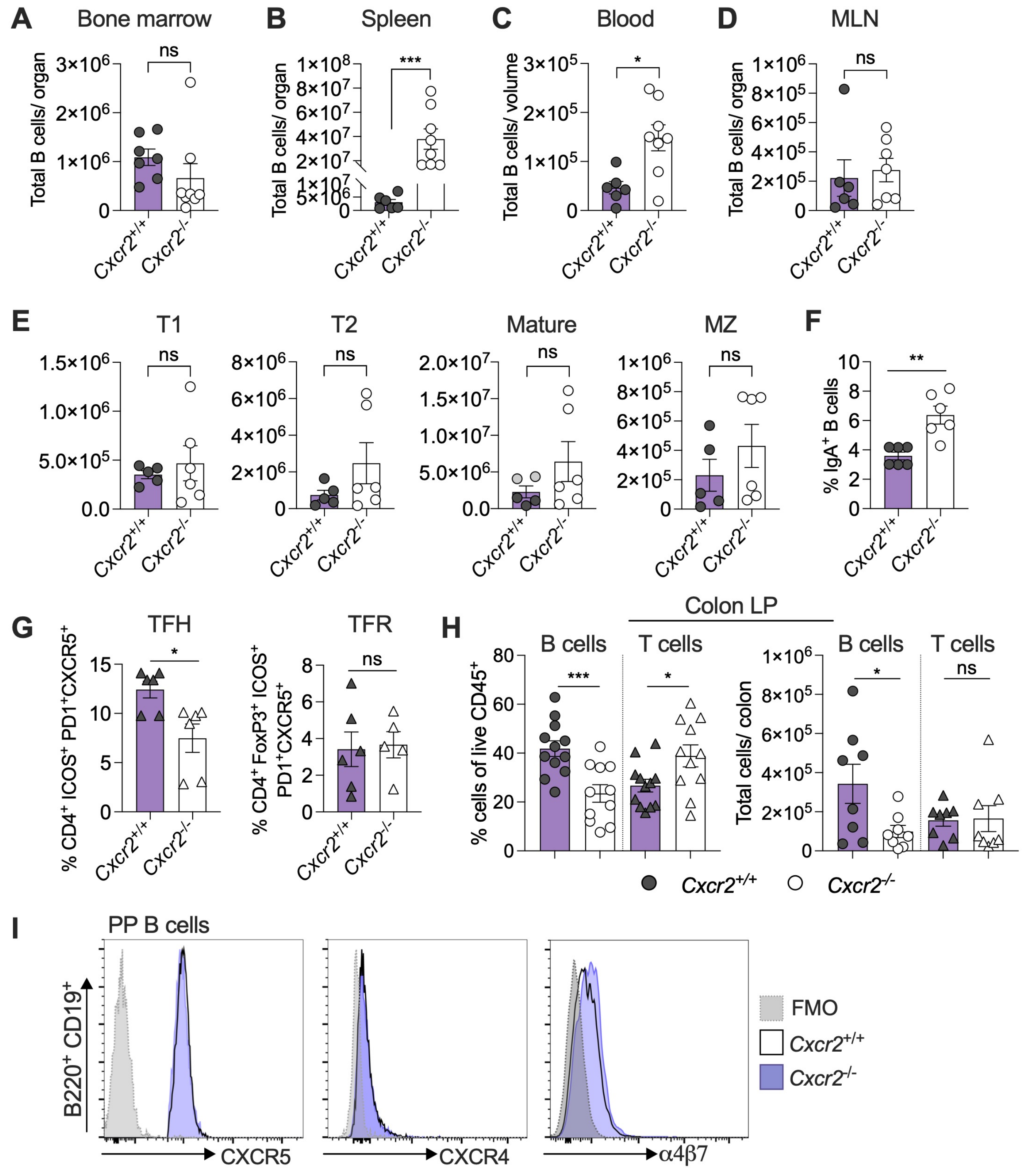


**fig. S3. Comparison of selected B and T cell populations in WT and *Cxcr2$^{-/-}$* mice.**
(**A – D**) Total CD19$^+$B220$^+$ B cell populations were quantified in the bone marrow, spleen, peripheral blood (volume = 300 μL), and mesenteric lymph nodes (MLNs) of *Cxcr2$^{+/+}$* and *Cxcr2$^{-/-}$* mice. **(E)** B cell developmental subsets in the spleen, including transitional B cells: IgM$^{hi}$IgD$^-$CD21$^-$ T1 B cells, IgM$^{hi}$IgD$^{hi}$CD21$^{hi}$CD23$^+$ T2 cells, IgM$^{hi}$IgD$^-$CD21$^{hi}$CD23$^-$ marginal zone (MZ) B cells, and IgM$^{low}$IgD$^{hi}$CD23$^+$ mature B cells were analyzed by flow cytometry (*15*). **(F)** The

frequency of IgA$^+$ B cells in Peyer's patches (PP) is shown. (**G**) Frequencies of T follicular helper (TFH) and T follicular regulatory (TFR) cells in PP were determined in *Cxcr2*$^{+/+}$ and *Cxcr2*$^{-/-}$ mice. (**H**) Frequencies and absolute numbers of CD19$^+$B220$^+$ B cells and CD3$^+$ T cells in the colonic lamina propria (LP) were determined by flow cytometry. (**I**) Expression levels of CXCR5, CXCR4, and α4β7 on *Cxcr2*$^{+/+}$ and *Cxcr2*$^{-/-}$ B cells are shown. FMO, fluorescence minus one. Bars represent the mean ± SEM. Significant differences are indicated by *p ≤0.05, **p<0.01, ***p<0.001, ns = not significant. Unpaired Student's *t* test.

**Supplemental Figure 4**

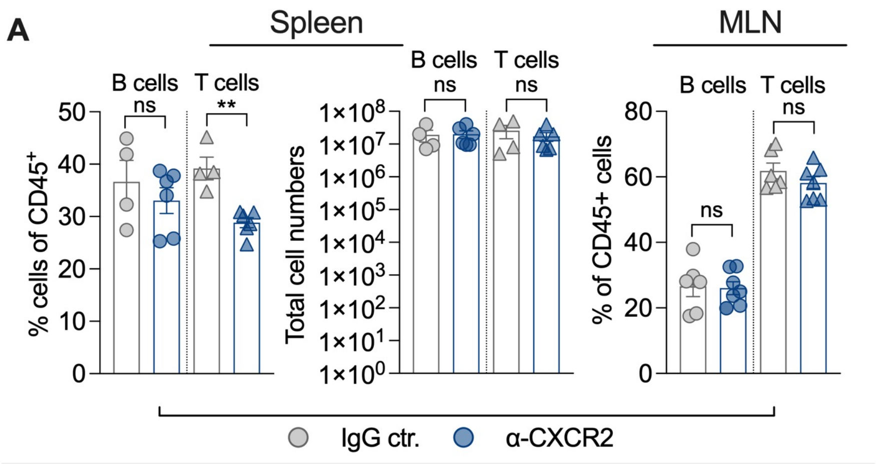


**fig. S4. B and T cell populations in the spleen and mesenteric lymph nodes of mice treated with CXCR2 antiserum.**

Frequencies and absolute numbers of CD19$^+$B220$^+$B cells and CD3$^+$ T cells in the spleen and mesenteric lymph nodes (MLN) of mice treated with CXCR2 antiserum or isotype-matched IgG control serum. Bars represent the mean ± SEM. Significant differences are indicated by **p<0.01, ns = not significant. Unpaired Student's *t* test.

**Supplemental Figure 5**

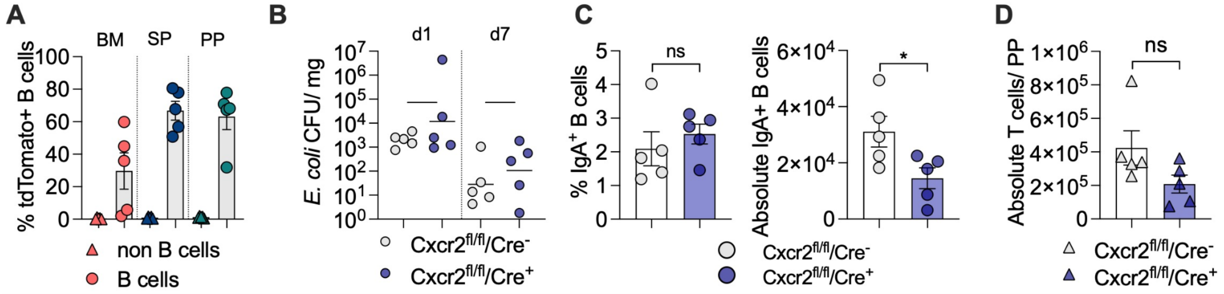


**fig. S5. Cell composition and *E. coli* colonization in mice with a targeted deletion of *Cxcr2* in B cells.**

(**A**) hCD20Tam-Cre x ROSA26$^{tdTomato}$ reporter mice were treated with tamoxifen for four consecutive days, followed by a 7-day rest period. Mice then received one more injection of tamoxifen, then after a 7-day rest period the frequencies of tdTomato$^+$ cells among CD19$^+$B220$^+$ B cells and CD19$^-$B220$^-$ cells were assessed in the bone marrow (BM), spleen (SP), and Peyer's patches (PP). (**B**) Fecal colony-forming units (CFU) of *E. coli* per mg feces on days 1 and 7 of the experiment in the respective experimental groups. (**C**) Frequencies and absolute numbers of $CD19^+B220^+IgA^+$ B cells in PP of respective groups as determined by flow cytometry. (**D**) Absolute numbers of $CD3^+$ T cells in Peyer`s patches (PP) of *Cxcr2*$^{fl/fl}$ x hCD20-TamCre$^+$ and *Cxcr2*$^{fl/fl}$ x hCD20-TamCre$^-$ mice were quantified by flow cytometry at the experimental endpoint. Bars represent the mean ± SEM (A, C, D) or geometric mean (B). Significant differences are indicated by *p ≤0.05, ns = not significant. Unpaired Student's *t* test for normally distributed groups or Mann-Whitney U for unpaired data sets that are not normally distributed.

**Table S1. Strains used in this study**

| Designation | Genotype | Reference or Source |
|---|---|---|
| EcN | *E. coli* Nissle 1917, wild type | Manuela Raffatellu (originally from Ardeypharm, Germany) |
| EcN Δ*csgA* | *E. coli* Nissle 1917 Δ*csgA*:: kan (Kan$^R$) | Cagla Tukel, (*13*) |
| *E. coli* RRG1 | Mouse commensal *E. coli* | This study |
| *B. theta* | *Bacteroides thetaiotamicron* strain VPI 5482 (ATCC 29148) | Hiutung Chu |
| *Clostridium* sp. 7_2 | *Clostridium* sp. 7_2 (HMP, #254) | Hiutung Chu |
| *E. cloacae* | *Enterobacter cloacae* subsp. *cloacae* strain CDC 442-68 (ATCC 13047) | Hiutung Chu |
| *Proteus mirabilis* | *Proteus mirabilis* BL95 | Judith Behnsen, (*14*) |
| *Lacticaseibacillus rhamnosus* | *Lacticaseibacillus rhamnosus* (HMP, #133) | Hiutung Chu |

**Table S2. Antibodies used in this study**

| Antigen | Vendor | clone | Application |
|---|---|---|---|
| Zombie Yellow Fixable Viability Kit | BioLegend | | FC |
| CD45R/B220 | BioLegend | RA3-6B2 | FC, IF |
| CD19 | BioLegend | 6D5 | FC |
| CD3 | eBioscience | 17A2 | FC |
| CD4 | eBioscience | RM4-5 | FC |
| CD8a | BioLegend | 53-6.7 | FC |
| CD45 | BioLegend | 30-F11 | FC |
| Ly6G | BioLegend | 1A8 | FC |
| CD11b | BioLegend | M1/70 | FC |
| IgM | BioLegend | RMM-1 | FC |
| F4/80 | BioLegend | BM8 | FC |

| | | | |
|---|---|---|---|
| IgD | BioLegend | 11-26c.2a | FC |
| IgA | eBioscience | mA-6E1 | FC |
| CD45.1 | BioLegend | A20 | FC |
| CD45.2 | BioLegend | 104 | FC |
| CD23 | BioLegend | B3B4 | FC |
| FoxP3 | BioLegend | 7E9 | FC |
| Integrin α4β7 | eBioscience | DATK-32 | FC |
| CD184/CXCR4 | BioLegend | L276F12 | FC |
| CD185/CXCR5 | BioLegend | L138D7 | FC |
| CD182/CXCR2 | BioLegend | SA044G4 | FC |
| CD16/32 (Fc block) | BioLegend | 93 | FC |
| CD3ε | BioLegend | 500A2 | IF |
| CXCR2 antiserum | Bio-Synthesis | polyclonal | In-vivo treatment |
| Rabbit gamma globulin | Jackson ImmunoResearch | polyclonal | In-vivo treatment |

Abbreviations: FC, flow cytometry; IF, immunofluorescence; Antibody clones were used in various fluorophore-conjugated formats.
